# Measuring SARS-CoV-2 neutralizing antibody activity using pseudotyped and chimeric viruses

**DOI:** 10.1101/2020.06.08.140871

**Authors:** Fabian Schmidt, Yiska Weisblum, Frauke Muecksch, Hans-Heinrich Hoffmann, Eleftherios Michailidis, Julio C. C. Lorenzi, Pilar Mendoza, Magdalena Rutkowska, Eva Bednarski, Christian Gaebler, Marianna Agudelo, Alice Cho, Zijun Wang, Anna Gazumyan, Melissa Cipolla, Marina Caskey, Davide F. Robbiani, Michel C. Nussenzweig, Charles M. Rice, Theodora Hatziioannou, Paul D. Bieniasz

## Abstract

The emergence of SARS-CoV-2 and the ensuing explosive epidemic of COVID19 disease has generated a need for assays to rapidly and conveniently measure the antiviral activity of SARS-CoV-2-specific antibodies. Here, we describe a collection of approaches based on SARS-CoV-2 spike-pseudotyped, single-cycle, replication-defective human immunodeficiency virus type-1 (HIV-1) and vesicular stomatitis virus (VSV), as well as a replication-competent VSV/SARS-CoV-2 chimeric virus. While each surrogate virus exhibited subtle differences in the sensitivity with which neutralizing activity was detected, the neutralizing activity of both convalescent plasma and human monoclonal antibodies measured using each virus correlated quantitatively with neutralizing activity measured using an authentic SARS-CoV-2 neutralization assay. The assays described herein are adaptable to high throughput and are useful tools in the evaluation of serologic immunity conferred by vaccination or prior SARS-CoV-2 infection, as well as the potency of convalescent plasma or human monoclonal antibodies.

## Introduction

The emergence of a new human coronavirus, SARS-CoV-2, in late 2019 has sparked an explosive global pandemic of COVID19 disease, with many millions of infections and hundreds of thousands of deaths (as of early June, 2020). The socioeconomic impact of the COVID19 pandemic has also been profound, with the mobility and productivity of a large fraction of the world’s population dramatically curtailed.

Human coronaviruses, including SARS-CoV-2, the other severe epidemic coronaviruses (MERS-CoV, SARS-CoV), and the mild the seasonal coronaviruses, all elicit neutralizing antibodies (Kellam and Barclay, 2020). These antibodies likely provide at least some degree of protection against reinfection. However, in the case of the seasonal coronaviruses, epidemiological and human challenge experiments indicate that protection is incomplete and diminishes with time, concurrent with declining neutralizing antibody titers (Callow et al., 1990; Kiyuka et al., 2018). The neutralizing antibody response to MERS-CoV and SARS-CoV is highly variable (Alshukairi et al., 2016; Cao et al., 2007; Choe et al., 2017; Liu et al., 2006; Mo et al., 2006; Okba et al., 2019; Payne et al., 2016), and because human infection by these viruses is rare (MERS-CoV) or apparently absent (SARS-CoV), the extent to which prior infection elicits durable protection against reinfection is unknown. For SARS-CoV-2, early studies, including our own, indicate that the magnitude of antibody responses is extremely variable, and a significant fraction of convalescents have comparatively low to undetectable levels of plasma neutralizing antibodies (Robbiani et al., 2020; Wu et al., 2020a). Thus, the effectiveness and durability of immunity conferred by primary SARS-CoV-2 infection is unknown, particularly in those who mount weaker immune response and is obviously a pressing issue, given the global spread of this virus. Moreover, because treatment and prevention modalities for SARS-CoV-2 are urgently sought, convalescent plasma is being evaluated for COVID19 therapy and prophylaxis (Bloch et al., 2020). Clearly, the effectiveness of such an intervention is likely to be profoundly impacted by the levels of neutralizing antibodies in donated convalescent plasma.

Effective vaccination and administration of cloned human monoclonal antibodies may be more successful than prior natural infection and convalescent plasma in providing antibody-based protection from SARS-CoV-2 infection. Indeed, recent work from our own laboratories and others has shown that closely related, highly potent, neutralizing monoclonal antibodies targeting the SARS-CoV-2 receptor binding domain (RBD) can be isolated from multiple convalescent donors (Brouwer et al., 2020; Cao et al., 2020; Chen et al., 2020b; Chi et al., 2020; Ju et al., 2020; Robbiani et al., 2020; Rogers et al., 2020; Seydoux et al., 2020; Shi et al., 2020; Wec et al., 2020; Wu et al., 2020b; Zost et al., 2020). Potent antibodies can be isolated from individuals with high or unexceptional plasma neutralizing titers, suggesting that natural infection in some individuals does not induce sufficient B-cell expansion and maturation to generate high levels of such antibodies (Robbiani et al., 2020; Wu et al., 2020a). However, these findings suggest that such antibodies might be straightforwardly elicited by vaccination.

Whether elicited by natural infection or vaccination, or administered as convalescent plasma or in recombinant form, neutralizing antibodies will likely be crucial for curtailing the global burden of COVID19 disease. For this reason, the availability of rapid, convenient and accurate assays that measure neutralizing antibody activity is crucial for evaluating naturally acquired or artificially induced immunity. Measuring SARS-CoV-2 neutralizing antibodies using traditional plaque reduction neutralization tests (PRNT) is labor intensive, requires biosafety level (BSL)-3 laboratory facilites and is not amenable to high throughput. Thus, various assays based on vesicular stomatitis virus (VSV) or human immunodeficiency virus type -1 (HIV-1) virions, pseudotyped with the trimeric SARS-CoV-2 spike protein that are high-throughput and can executed at BSL-2 will be essential to evaluate neutralization activity. These pseudotype virus assays offer numerous advantages (Crawford et al., 2020; Nie et al., 2020), but their ability to predict plasma neutralization activity against authentic SARS-CoV-2, or correctly identify the most potent human monoclonal antibodies has not been rigorously evaluated.

Herein, we describe assays based on pseudotyped and chimeric viruses that our laboratories have used to measure the neutralizing activity of convalescent plasma and to identify potently neutralizing human monoclonal antibodies against SARS-CoV-2. These assays are rapid and convenient. Using a panel of convalescent plasma and human RBD-specific monoclonal antibodies, we demonstrate that these assays provide measurements of virus neutralization that are well correlated with a neutralizing antibody test employing authentic SARS-CoV-2 virions. As such, these tools are useful to estimate SARS-CoV-2 immunity in the context of recovery from infection, in experimental vaccine recipients and to evaluate the potency of antibody-based therapy and prophylaxis

## Results

### HIV-1-based SARS-CoV-2 S pseudotyped virions

To generate SARS-CoV-2 pseudotyped HIV-1 particles, we constructed a replication defective HIV-1 proviral plasmid (pHIV-1_NL_ΔEnv-NanoLuc Fig. 1A) that lacks a functional viral *env* gene, and contains sequences encoding a NanoLuc luciferase protein in place of the *nef* gene. This proviral construct is similar to the widely used pNL4-3.Luc.R-E-proviral reporter plasmid (Connor et al., 1995), but NanoLuc luciferase yields ∼100-fold brighter luminescence than firefly luciferase, facilitating the detection of small numbers of infected cells. Indeed, we estimate that single infection events can be detected in a 96-well assay format (see below).

**Fig. 1.**
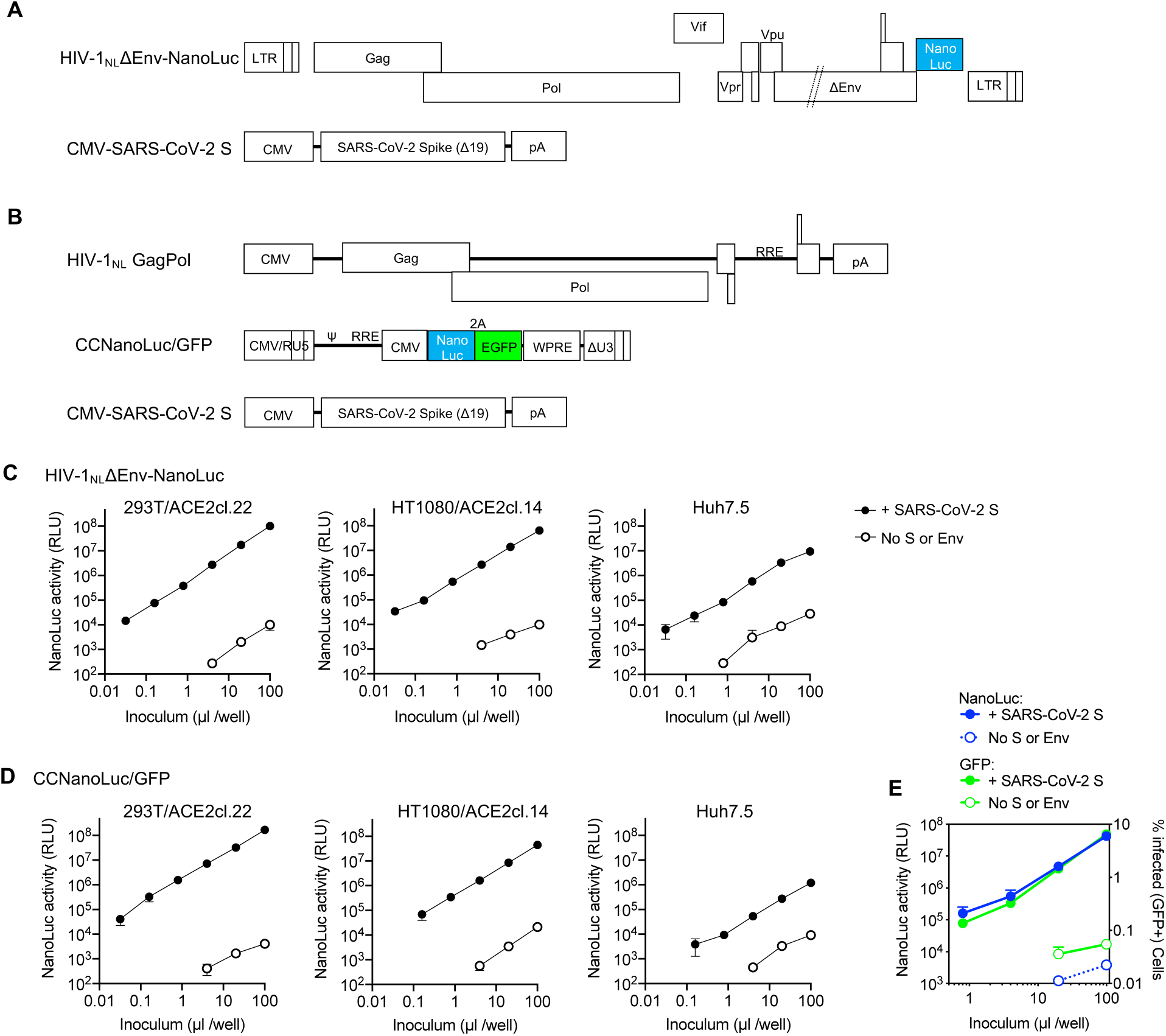
Two-plasmid and three-plasmid HIV-1-based pseudotyped viruses. **A**. Schematic representation of the modified HIV-1_NL_ ΔEnv-NanoLuc genome in which a deletion in *env* was introduced and Nef-coding sequences were replaced by those encoding a NanoLuc luciferase reporter. Infectious virus particles were generated by cotransfection of pHIV-1_NL4_ΔEnv-NanoLuc and a plasmid encoding the SARS-CoV-2 S lacking the 19 amino acids at the C-terminus of the cytoplasmic tail (SΔ19). **B**. Schematic representation of constructs used to generate SARS-CoV-2 S pseudotyped HIV-1-based particles in which HIV-1_NL_GagPol, an HIV-1 reporter vector (pCCNanoLuc/GFP) encoding both NanoLuc luciferase and EGFP reporter and the SARS-CoV-2 SΔ19 are each expressed on separate plasmids. **C**. Infectivity measurements of HIV-1_NL_ ΔEnv-NanoLuc particles (generated using the plasmids depicted in A) on the indicated cell lines. Infectivity was quantified by measuring NanoLuc luciferase activity (Relative Light Units, RLU) following infection of cells in 96-well plates with the indicated volumes of pseudotyped viruses. The mean and standard deviation of two technical replicates is shown. Target cells 293T/ACE2cl.22 and HT1080/ACE2cl.14 are single-cell clones engineered to express human ACE2 (see Fig S1A). Virus particles generated in the absence of viral envelope glycoproteins were used as background controls. **D**. Same as, C but viruses were generated using the 3 plasmids depicted in B. **E**. Infectivity meaurements of CCNanoLuc/GFP containing SARS-CoV-2 pseudotyped particles generated using plasmids depicted in B on 293ACE2*(B) cells, quantified by measuring NanoLuc luciferase activity (RLU) or GFP levels (% of GFP positive cells). Mean and standard deviation from two technical replicates is shown.

Because some localities require that two-plasmid HIV-1-based pseudotyped viruses be used in BSL2+ or BSL3 laboratories, we also developed HIV-1 pseudotyped viruses using a three-plasmid approach in which the packaged viral-vector genome and GagPol expression functions are installed on separate plasmids (Fig. 1B). We also constructed a packageable HIV-1 vector (pCCNanoLuc/GFP) that encodes both a NanoLuc luciferase reporter and a GFP reporter (Fig. 1B). This 3-plasmid format recapitulates that used in commonly used lentivirus vector procedures, except that the conventionally used VSV-G envelope expression plasmid is omitted. Instead, for both the two-plasmid (HIV-1_NL_ΔEnv-NanoLuc) and three-plasmid (CCNanoLuc/GFP) HIV-1 pseudotype formats, we constructed plasmids encoding codon-optimized SARS-CoV-2 spike (S) proteins (Fig. 1A, B). We also generated several 293T and HT1080 derived cell lines expressing the SARS-CoV and SARS-CoV-2 receptor ACE2 (Li et al., 2003), of which several populations and clones expressing varying levels of ACE2 were used herein (Fig. S1 A,B).

Incorporation of envelope or spike proteins into heterologous viral particles is unpredictable. Indeed, even minor alterations to the cytoplasmic tail of the HIV-1 Env protein can block its incorporation into homologous HIV-1 particles (Murakami and Freed, 2000). For this reason, we compared infection using CCNanoLuc/GFP particles pseudodotyped with either the full-length SARS-CoV-2 spike protein, or derivatives with either 18 or 19 amino acids truncated from the C-terminus. While the full-length SARS-CoV-2 S protein supported the generation of infectious virions that gave a luminescence signal higher than that of virions lacking S, both the Δ18 and Δ19 truncated forms generated ∼10-fold higher titers of infectious particles than the full-length SARS-CoV-2 S protein (Fig. S1C). Thus, the Δ19 variant of SARS-CoV-2 was used hereafter unless otherwise indicated. Similarly, the SARS-CoV S protein generated infectious virions, but higher infectious titers were obtained when a Δ18 and Δ19 truncated SARS-CoV S protein was used (Fig. S1C).

Both the two-plasmid (HIV-1_NL_ΔEnv-NanoLuc) and three-plasmid (pCCNanoLuc/GFP) derived SARS-CoV-2 pseudotyped viruses infected ACE2-expressing 293T and HT1080 cells, yielding a strong luminescence signal of up to 10^7^ to10^8^ relative light units (RLU), while unmanipulated parental cell lines were infected poorly (293T), or not at all (HT1080) (Fig 1C, D, Fig S1D, E). A cell line (Huh7.5) that endogenously expresses ACE2 was also infected by both HIV-1 SARS-CoV-2 pseudotypes, although the luminescent signal was not as high as in the engineered 293T/ACE2 or HT1080/ACE2 cell lines (Fig. 1 C, D). The SARS-CoV-2 pseudotyped HIV-1 virions could be concentrated by ultracentrifugation and without loss of titer and without effects on the background level of NanoLuc luciferase (Fig. S1F).

Examination of the panel of ACE2-expressing 293T derived cell lines revealed that the level of infection appeared dependent on the level of ACE2 expression (Fig S1A, S2A, S2B). Conversely, the level of infection was not dramatically affected by varying the amount of co-transfected pSARS-CoV-2-S_Δ19_ during pseudotyped virus production (Fig S2C).

A common misconception is that the maximum dynamic range of infection assays measured via expression of a viral genome-encoded luciferase reporter (such as those described herein) is the same as the dynamic range of the luciferase assay. The ability to count the number of CCNanoLuc/GFP infected cells (by flow cytometry) and measure NanoLuc luciferase activity in replicate wells (Fig. 1E), afforded the ability to readily determine the average NanoLuc luciferase activity generated by a single CCNanoLuc/GFP infectious unit. This calculation led to the conclusion that a single infectious unit of CNanoLuc/GFP pseudotyped virus generated an average of approximately 1.2 × 10^4^ RLU in infected 293T/ACE2*(B) cells. A second estimate, based on immunofluorescent detection of HIV-1 Gag expressed in 293T/ACE2(B) cells following infection with the HIV-1_NL_ΔEnv-NanoLuc/SARS-CoV-2 pseudotype (Fig. S2D) suggested that single infected cells generate approximately 5 × 10^3^ RLU. Given that the highest signals generated in the NanoLuc luciferase assays following infection with SARS-CoV-2 pseudotyped HIV-1_NL_ΔEnv-NanoLuc or CNanoLuc/GFP are between 10^7^ and 10^8^ RLU (depending on the target cell line, Fig 1C, D, Fig S1C-E), this value is commensurate with the observation that ∼10% or greater of the 10^4^ cells plated in each well became infected (Fig. S2A). Thus, the dynamic range of these HIV-1 pseudotype infection assays, formatted in 96-well plates, is between 3 and 4 orders of magnitude, depending on the amount of pseudotyped virus and the particular target cell line used. This dynamic range is more than adequate for accurate determinations of plasma neutralizing activity as well as determination of potency (IC_50_ and IC_90_) of monoclonal antibodies or other inhibitors of SARS-CoV-2 S-dependent viral entry.

### VSV-based SARS-CoV-2 S pseudotyped virions

Another commonly used platform for evaluation of virus envelope or spike protein function is based on vesicular stomatitis virus (VSV) lacking a G protein (VSVΔG) (Whitt, 2010). This approach is possible because VSVΔG can replicate well when complemented *in trans* by either its own envelope (G) protein or (sometimes) by a heterologous viral glycoprotein. We constructed a VSVΔG genome that contained a dual reporter (mNeonGreen and NanoLuc luciferase) termed rVSVΔG/NG-NanoLuc (Fig. 2A). The dual reporter enabled infection by rVSVΔG/NG-NanoLuc pseudotype to be monitored by imaging (Fig S3A), flow cytometry or NanoLuc luciferase assay (Fig. 2B). An advantage of VSV pseudotypes over their HIV-1 counterparts is that the rapid intracellular replication of the VSV genome enables robust reporter gene expression to be detected within a few hours after infection (Fig 2C Fig S3B). To maximize signal over background, we used an overnight 16h infection protocol, unless otherwise stated. Like HIV-1 pseudotypes, the rVSVΔG/NG-NanoLuc pseudotypes selectively infected ACE2 expressing 293T and HT1080 cells, although unmodified 293T cells also exhibited low level susceptibility (Fig. 2D, Fig. S3C, D). As was the case with HIV-1 pseudotypes, the level of rVSVΔG/NG-NanoLuc/SARS-CoV-2 infection was dependent on the level of ACE2 expression (Fig. S3E), although the levels of endogenously expressed ACE2 in Huh7.5 cells and Vero E6 cells were sufficient to give a robust signal (Fig. 2D). Indeed, we used Huh7.5 cells hereafter, unless otherwise indicated

**Fig. 2.**
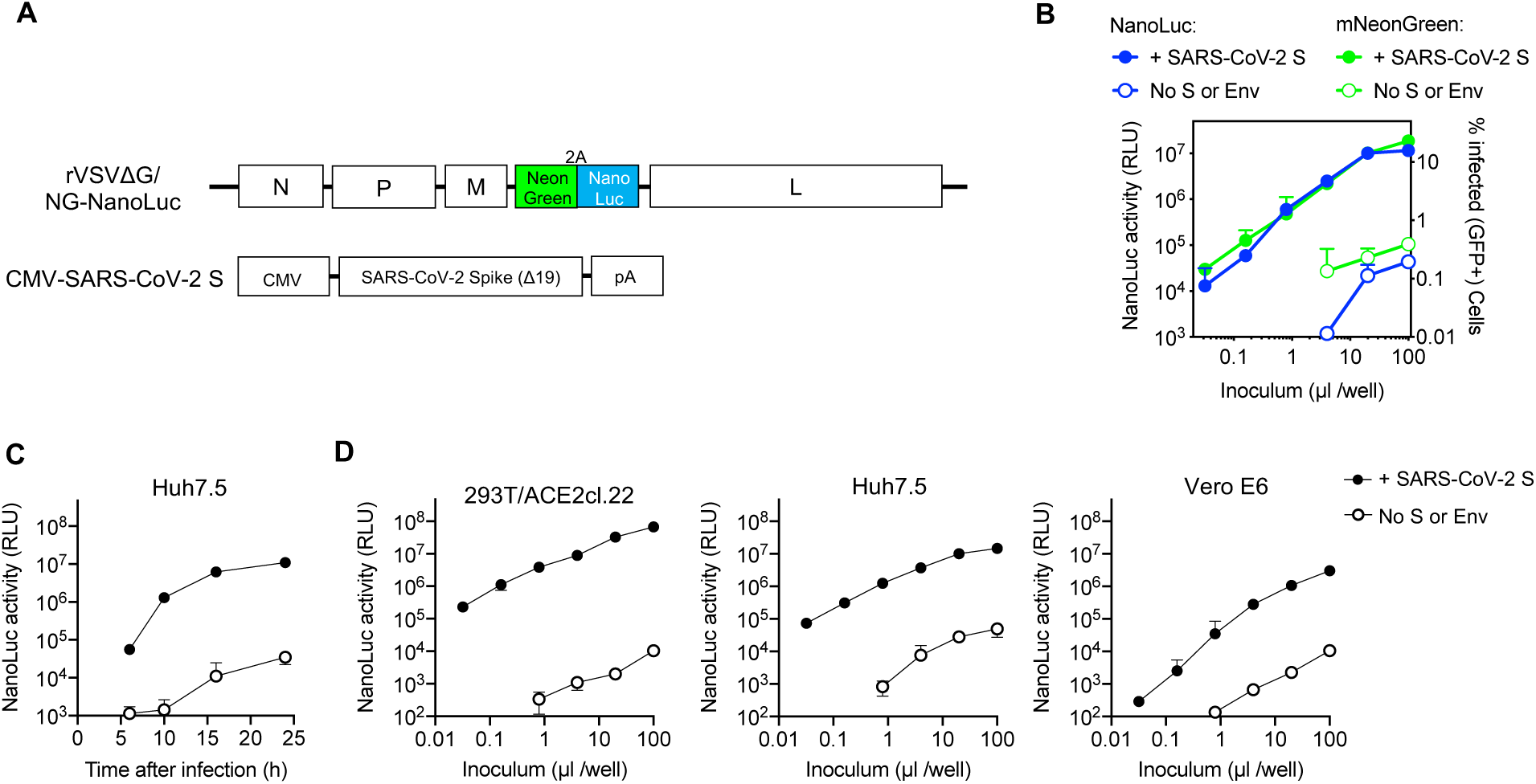
VSV-based SARS-CoV-2 pseudotyped viruses. **A**. Schematic representation of the rVSVΔG/NG-NanoLuc genome in which G-coding sequences were replaced by an mNeonGreen-2A-NanoLuc luciferase reporter cassette. Infectious virus particles were generated by passaging G-complemented rVSVΔG/NG-NanoLuc virus stocks through 293T cells transfected with a plasmid encoding SARS-CoV-2 SΔ19. **B**. Infectivity of pseudotyped rVSVΔG/NG-NanoLuc particles on Huh7.5 cells was quantified by measuring luciferase activity (RLU) or the % GFP-positive cells. Mean and standard deviation from two technical replicates is plotted. Virus particles generated by passage through cells that were not transfected with SARS-CoV-2 S were used as a control. **C**. NanoLuc luciferase activity (RLU) in Huh7.5 cells measured at various times after infection with pseudotyped rVSVΔG/NG-NanoLuc particles. Average and standard deviation from two technical replicates is shown. **D**. Infectivity of pseudotyped rVSVΔG/NG-NanoLuc particles on the indicated cell lines. Infectivity was quantified by measuring NanoLuc luciferase activity (RLU) following infection of cells in 96-well plates with the indicated volumes of pseudotyped viruses. Average and standard deviation from two technical replicates is shown.

A disadvantage of the NanoLuc luciferase reporter, is that this protein is highly stable, more so than other luciferases. Because rVSVΔG replication is quite cytopathic, pseudotype virion preparations were contaminated with NanoLuc luciferase protein, which elevated the assay background. However, this problem could be relieved by pelleting virions by ultracentrifugation through sucrose, or concentration using Lenti-X (Fig S3F). Indeed, counting of mNeonGreen infected cells and luciferase quantitation (Fig 2B, Fig S3A) revealed that individual infected Huh7.5 cells yielded ∼6×10^3^ RLU per infected cell. Thus, like the HIV-1-based assay, the rVSVΔG/NG-NanoLuc/SARS-CoV-2 pseudotype infection assay had a dynamic range of 3 to 4 orders of magnitude, when formatted in 96-well plates.

### Construction of a replication competent VSV/SARS-CoV2 chimeric virus

The aforementioned assays both employ single-cycle, replication-defective constructs and do not allow for any viral spread in the presence of antibody. This feature could impact the sensitivity with which neutralizing activity is detected (see discussion). To construct a replication competent VSV/SARS-CoV-2 chimera, we inserted sequences encoding the SARS-CoV-2 S protein lacking the C-terminal 18 codons into a recombinant vesicular stomatitis virus (VSV) background that contains green fluorescent protein (GFP) cDNA between the inserted S sequence and the L (polymerase) gene. Thus, in this construct, the SARS-CoV-2 S replaces the native VSV-G protein (Fig. 3A). The recombinant virus was rescued in 293T cells by co-expressing T7 polymerase and complementing VSV proteins. The rescued virus was passaged once in 293T cells transfected with a VSV-G-expression plasmid to facilitate initial virus amplification. Since G is not encoded in the recombinant virus genome, subsequent rounds of infection were dependent on the SARS-CoV-2 S protein. Thus, thereafter the complemented rVSV/SARS-CoV-2/GFP virus was used to infect 293T/ACE2(B) cells (in the absence of the complementing VSV-G protein) and infection monitored by observation of GFP expression. Initially, the rescued rVSV/SARS-CoV-2 replicated poorly – several days were required for the majority of the cells in the culture to become infected (GFP-positive). Supernatant (500 μl/75cm^2^ flask) from these infected 293T/ACE2(B) cells was used to infect fresh 293T/ACE2(B) cells, and this process was repeated for three additional passages. After three passages, accelerated replication was clearly observed. Specifically, all cells in a 75cm^2^ flask became GFP-positive within 24 h of inoculation with 100 μl of supernatant from the previous passage. Thereafter, individual viral variants were isolated by limiting dilution in 96-well plates containing 293T/ACE2(B) cells. Cells and supernatant were harvested from two wells that each contained one individual GFP-positive plaque (signifying infection by single viruses) and designated rVSV/SARS-CoV-2/GFP_1D7_ and rVSV/SARS-CoV-2/GFP_2E1_.

**Fig. 3.**
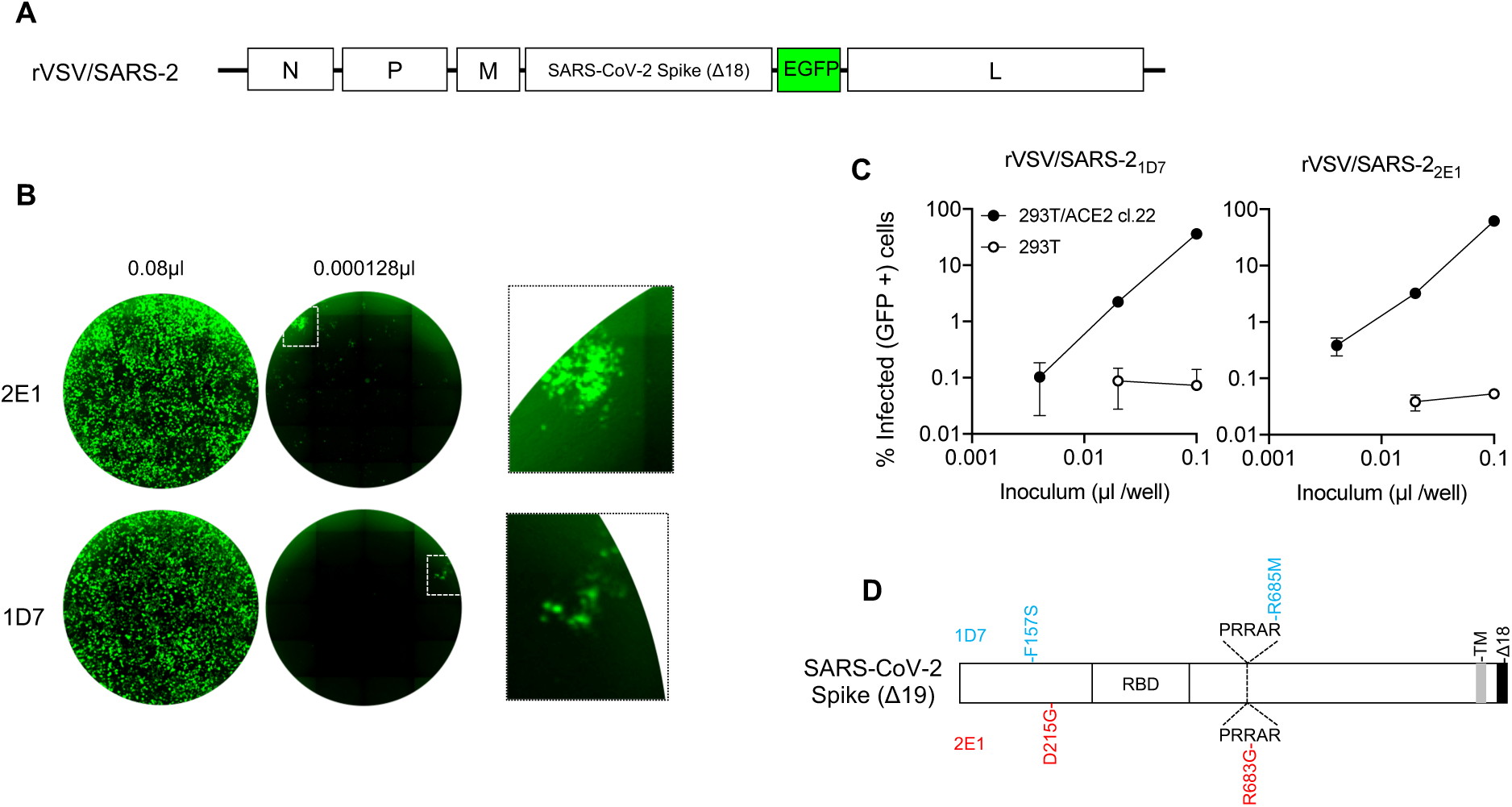
A replication-competent VSV/SARS-CoV-2 chimera. **A**. Schematic representation of the rVSV/SARS-CoV-2/GFP genome in which G-encoding sequences were replaced by SARS-CoV-2 SΔ18 coding sequences. GFP-encoding sequences were introduced between the SARS-CoV-2 SΔ18 and L open reading frames. **B**. Representative images of 293T/ACE2(B) cells infected with the indicated volumes of plaque purified, adapted derivatives (2E1 and 1D7) of VSV/SARS-CoV-2/GFP following passage in the same cell line. Left and center images show contents of an entire well of a 96-well plate, the right image shows expanded view of the boxed areas containing individual plaques. **C**. Infectivity measurements of rVSV/SARS-CoV-2/GFP virus stocks on 293T/ACE2(B) or control 293T cells, quantified by measuring % GFP-positive cells at 16h after infection. Average and standard deviation from two technical replicates is shown. **D**. Schematic representation of the adaptive changes acquired in rVSV/SARS-CoV-2/GFP during passage. Changes in 1D7 and 2E1 are shown in blue and red, respectively.

The adapted rVSV/SARS-CoV-2/GFP_1D7_ and rVSV/SARS-CoV-2/GFP_2E1_ both grew rapidly and achieved titers of between 1×10^7^ and 1×10^8^ plaque forming units/ml following replication for 36-48h in 293T/ACE2(B) or 293T/ACE2cl.22 cells. (Fig 3B, C). We extracted RNA from the supernatant of cells infected each of these viruses and determined the nucleotide sequence of the introduced SARS-CoV-2 spike cDNA and flanking regions. The S proteins were each found to encode two nonsynonymous changes: rVSV/SARS-CoV-2/GFP_1D7_ encoded F157S and R685M mutations, while rVSV/SARS-CoV-2/GFP_2E1_ encoded D215G and R683G mutations (Fig. 3D). Notably, R685M and R683G both alter the putative furin-like protease cleavage site in the SARS-CoV-2 S protein. Other isolated plaques whose SARS-CoV-2 S-encoding regions were sequenced, but were not further investigated, also contained furin cleavage site mutations (R682G or R685K) suggesting that modification of the furin cleavage site is a key adaptation for high-level rVSV/SARS-CoV-2 replication in 293T/ACE2 cells.

### Neutralization of pseudotyped HIV-1 and VSV, chimeric VSV/SARS-CoV-2 and authentic SARS-CoV-2 by antibodies

Because infection by the aforementioned viruses is dependent on SARS-CoV-2 S and ACE2 proteins, these assays should be good surrogates for the measurement of the SARS-CoV-2 neutralizing activity of convalescent plasma or candidate therapeutic/prophylactic monoclonal antibodies. Indeed, we have made extensive use of HIV-1_NL_ΔEnv-NanoLuc virions to measure levels of neutralizing activity in plasma of COVID19 patients, and to identify potent human monoclonal antibodies (Robbiani et al., 2020). Notably, the use of dual GFP and NanoLuc reporters in pseudotyped viral genomes provides for a rapid and flexible assessment of neutralizing activity, that can be assessed quantitively either microscopically or by flow cytometry and NanoLuc luciferase assays, which can be used interchangeably (Fig S4A, B, C).

To compare the neutralization properties of the aforementioned pseudotyped and chimeric viruses with authentic SARS-CoV-2, we used an antibody staining-based SARS-CoV-2 infection assay to quantify the numbers SARS-CoV-2 infected cells in 96-well plates. Twenty convalescent plasma samples that displayed a range of neutralization activities against authentic SARS-CoV-2 (Fig. 4A) were also evaluated in the SARS-CoV-2 pseudotyped HIV-1_NL_ΔEnv-NanoLuc and rVSVΔG/NG-NanoLuc as well as in replication competent VSV/SARS-CoV-2 neutralization assays (Fig 4B). Despite the fact that these neutralization assays have quite different dynamic ranges, employ different virion scaffolds and target cell lines, and involve single-cycle replication-defective or multi-cycle replication-competent viruses, the plasma neutralization titers obtained with the surrogate virus approaches were each well correlated with titers obtained using authentic SARS-CoV-2 (Fig. 4C). Notably, the HIV-1_NL_ΔEnv-NanoLuc and rVSVΔG/NG-NanoLuc pseudotyped viruses gave NT_50_ values that indicated marginally reduced sensitivity to plasma antibodies as compared to authentic SARS-CoV-2, with the rVSVΔG/NG-NanoLuc appearing to more sensitively detect weak plasma neutralizing activity (Fig 4C) than HIV-1_NL_ΔEnv-NanoLuc. Conversely, the replication competent VSV/SARS-CoV-2 was more sensitive to plasma neutralization than authentic SARS-CoV-2. Nevertheless, each of the surrogate viruses was able to provide a good indication of plasma neutralizing potency against authentic SARS-CoV-2

**Fig. 4.**
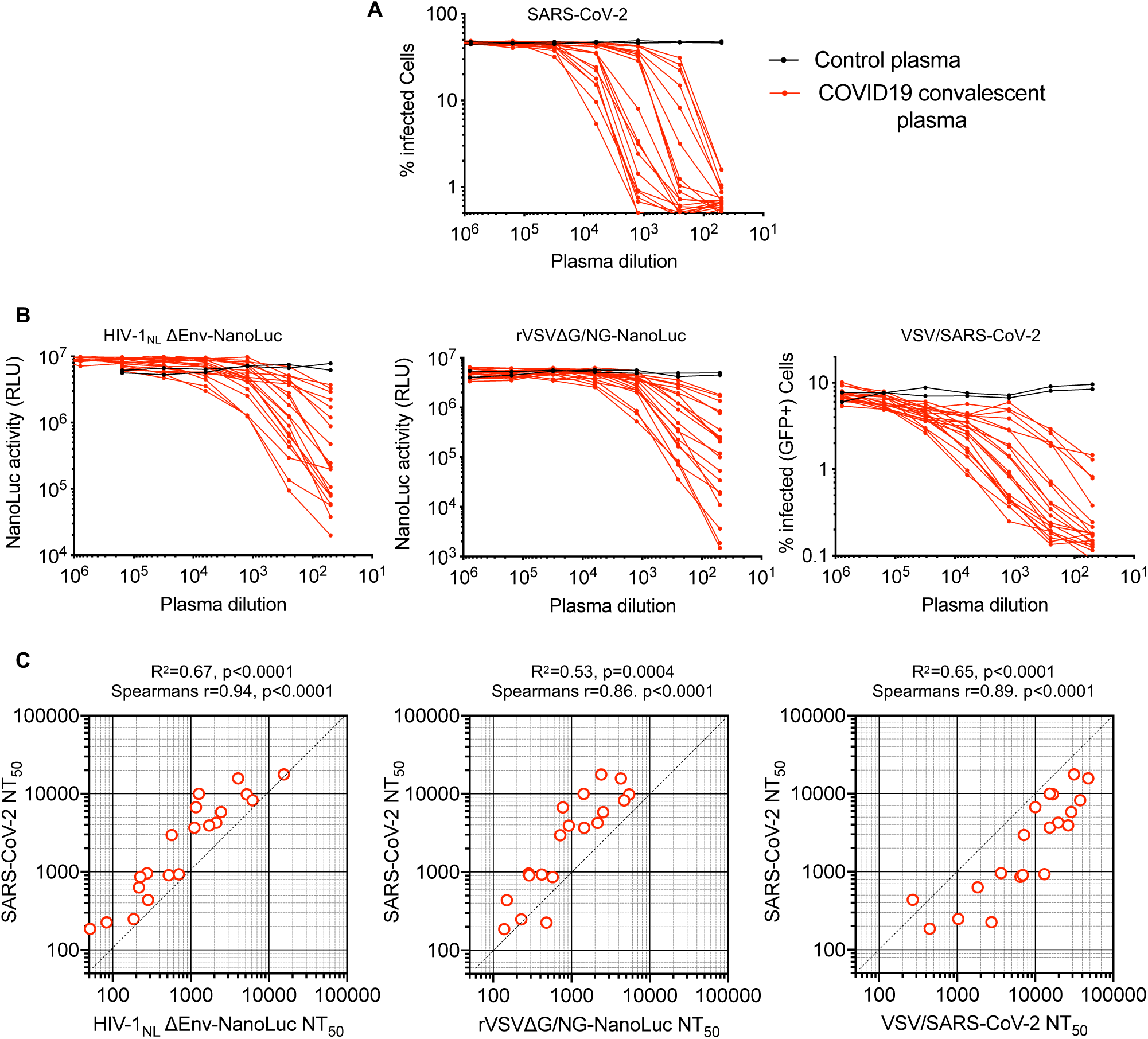
Measurement of neutralization activity in COVID19 convalescent donor plasma. **A**. Plasma neutralization of SARS-CoV-2: serial 5-fold dilutions of plasma samples from convalescent donors were incubated with SARS-CoV-2 n=3 replicates and residual infectivity determined using VeroE6 target cells, expressed as % infected cells by immunostaining. **B**. Plasma neutralization of HIV-1_NL_ΔEnv-NanoLuc pseudotyped virus using 293T/ACE2*(B) target cells, rVSVΔG/NG-NanoLuc pseudotyped virus using Huh7.5 target cells or replication competent rVSV/SARS-CoV-2/GFP using 293T/ACE2(B) target cells. Residual infectivity was quantified by measuring either NanoLuc luciferase (RLU) or the % GFP-positive cells, as indicated. **C**. Correlation between NT_50_ values for each of the 20 plasmas for each of the surrogate viruses (x-axis) and NT_50_ values for the same plasmas for SARS-CoV-2 (y-axis).

Next, we evaluated 15 human monoclonal antibodies that were identified by sorting of individual SARS-CoV-2 RBD-binding B-cells (Robbiani et al., 2020). The panel was selected on the basis of neutralization potency using the HIV-1_NL_ΔEnv-NanoLuc assay, and all had IC_50_ values ranging between 3ng/ml and 60ng/ml in this assay. These antibodies all potently neutralized SARS-CoV-2 (Fig 5A), as well as the HIV-1_NL_ΔEnv-NanoLuc and rVSVΔG/NG-NanoLuc pseudotyped viruses and the replication competent VSV/SARS-CoV-2 viruses (Fig 5B). Each of the surrogate viruses gave IC_50_ values for the antibody panel that correlated well with IC_50_ values measured using authentic SARS-CoV-2 (Fig. 5C). Interestingly, among the surrogate viruses, the two VSV-based viruses were the most different in terms of relative sensitivity to the monoclonal antibody panel. Specifically, while there was a good linear relationship between the IC_50_ values measured using SARS-CoV-2 and the rVSVΔG/NG-NanoLuc pseudotype virus, the latter was generally less sensitive to neutralization (Fig. 5C). The replication competent VSV/SARS-CoV-2/GFP virus appeared to most accurately predict the IC_50_ values of the monoclonal antibodies against authentic SARS-CoV-2, despite the fact that, unlike the pseudotyped viruses, its S protein encoded adaptive changes that arose during adaptation (Fig 3D). Importantly however, the most potent monoclonal antibodies had IC_50_ values of less than 10ng/ml, measured using all four viruses, indicating that each surrogate virus could correctly identify the most potently neutralizing human monoclonal antibodies.

**Fig. 5.**
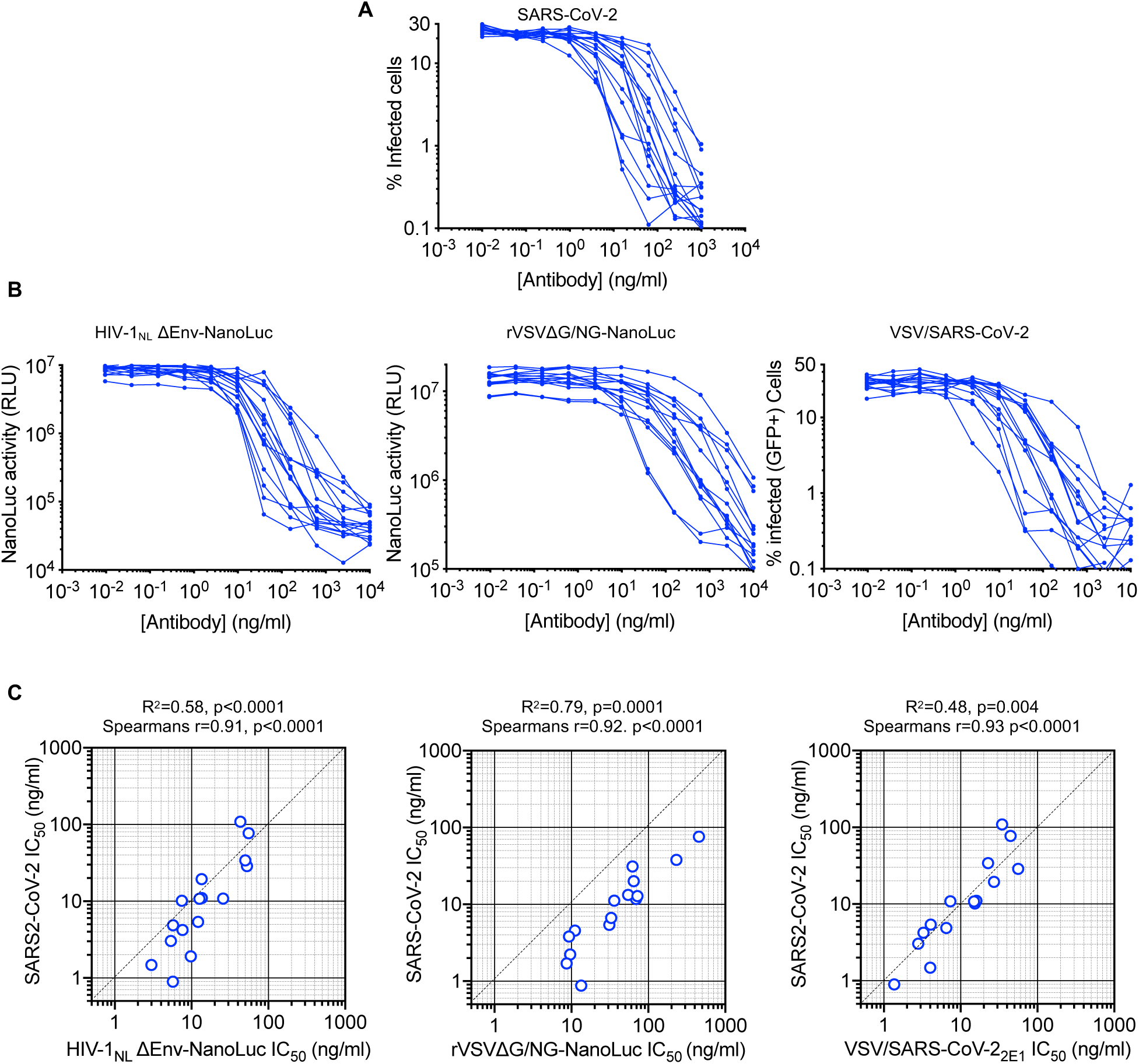
Measurement of neutralization potency of human monoclonal antibodies. **A**. Neutralization of SARS-CoV-2: the indicated concentrations of monoclonal antibodies were incubated with SARS-CoV-2 n=3 replicates and residual infectivity determined using Vero E6 target cells, expressed as % infected cells, by immunostaining **B**. Monoclonal antibody neutralization of HIV-1_NL_ΔEnv-NanoLuc pseudotyped virus using 293T/ACE2*(B) target cells, rVSVΔG/NG-NanoLuc pseudotyped virus using Huh7.5 target cells or replication competent rVSV/SARS-CoV-2/GFP using 293T/ACE2(B) target cells. Residual infectivity was quantified by measuring either NanoLuc luciferase (RLU) or the % GFP positive cells, as indicated. **C**. Correlation between IC_50_ values for each of the 15 monoclonal antibodies for each of the surrogate viruses (x-axis) and IC_50_ values for the same antibodies for SARS-CoV-2 (y-axis).

## Discussion

Herein, we describe pseudotyped and chimeric viruses that can evaluate the neutralizing activity of SARS-CoV-2 S-specific monoclonal antibodies as well as convalescent sera or plasma. Many factors could, in principle, affect the apparent potency of neutralizing antibodies as measured using surrogate viruses. One key factor may be the density of spikes on the virion envelope. Spike density could affect the avidity of bivalent antibodies, particularly those that are unable to engage two S-protein monomers within a single trimer and whose potency is dependent on engaging two adjacent trimers {Barnes, 2020 #41}(Galimidi et al., 2015). Electron microscopic images of coronaviruses indicate a fairly high spike density, and the extent to which pseudotyped viruses mimic this property might be important for determining the accuracy of neutralization assays. Notably, we found that truncation of the cytoplasmic tail of SARS-CoV-2 dramatically increased the infectious titer of SARS-CoV-2 pseudotypes, likely by facilitiating incorporation of S protein into virions. During virion assembly SARS-CoV-2 buds into secretory compartments (Stertz et al., 2007), while HIV-1 and VSV assemble at the cell surface (Jouvenet et al., 2006). Truncation of the S protein cytoplasmic tail may increase cell surface levels, and/or enable incorporation by alleviating structural incompatibility of the S protein cytoplasmic tail and HIV-1 or VSV matrix proteins.

To the extent that RBD-specific antibodies may compete with target cell surface ACE2 for binding to virion spikes, the density of ACE2 molecules on the target cell surface could additionally affect antibody potency in neutralization assays. Moreover, the use of replication competent, multicycle replication-based assays, versus single-cycle infection with defective reporter viruses could additionally affect apparent neutralizing antibody potency. Partial neutralization at marginal antibody concentrations in a single replication cycle might be amplified over multiple rounds of replication. Alternatively, the increase in viral dose during multiple replication cycles, or the high multiplicity associated with direct cell-to-cell viral spread might overwhelm neutralizing antibodies at marginal dilutions in multi-cycle neutralization assays. Such a scenario would reduce apparent antibody potency compared to single-cycle assays. Finally, viruses may also generate defective or noninfectious particles to varying degrees which could sequester neutralizing antibodies and therefore affect neutralization potency. Despite the very different nature of the assays employed herein, as well as their different dynamic ranges, each of the surrogate virus-based assays generated quantitative measurements of neutralizing activity that correlated well with neutralization measured using authentic SARS-CoV-2. Naturally, the above considerations mean that these correlations are not precise; for example, both HIV-1 and VSV-based pseudotyped viruses were somewhat less sensitive to neutralization than authentic SARS-CoV-2, particularly by weakly neutralizing plasma. This finding may be because they are single-cycle assays, or perhaps because pseudotyped virions may have lower spike density than SARS-CoV-2.

In addition to replication-defective single-cycle pseudotyped viruses we also developed a replication competent rVSV/SARS-CoV-2/GFP chimeric virus. Notably, adaptation of rVSV/SARS-CoV-2/GFP in 293T/ACE2 cells led to the acquisition of mutations at the S protein furin cleavage site. Adaptation of rVSV/SARS-CoV-2/GFP in different target cells, that express different furin-like or other proteases may result in the acquisition of alternative adaptive mutations. Crucially, the sensitivity of the adapted rVSV/SARS-CoV-2/GFP to neutralization by monoclonal antibodies mimicked that of authentic SARS-CoV-2. Interestingly, the rVSV/SARS-CoV-2/GFP appeared slightly more susceptible to plasma neutralization than SARS-CoV-2, for unknown reasons. Given that the design of rVSV/SARS-CoV-2/GFP is similar to that of the successful VSV/EboV ebolavirus vaccine, derivatives of the adapted 1D7 or 2E1 viruses could potentially be vaccine candidates. Additionally, these viruses could also be used in laboratory selection experiments to identify mutations that enable escape from inhibition by antibodies or other therapeutic agents that target the spike protein.

Some of the pseudotype neutralization assays described herein can be executed in a standard BSL-2 laboratory. Indeed, we have used the HIV-1 and VSV pseudotype approaches to conduct determine the neutralizing potencies of hundreds of plasma samples and monoclonal antibodies in a BSL-2 laboratory in a few weeks. Automation and additional miniaturization is certainly feasible to further increase throughput – a notable consideration given the sheer number of vaccine candidates in the development pipeline (Chen et al., 2020a). We note however, that miniaturization reduces the number of infected cells and the dynamic range of neutralization assays. We also note that the HT1080/ACE2cl.14 and Huh7.5 cell lines are significantly more adherent than 293T-derived cell lines and are recommended (for HIV-1 and VSV pseudotype assays, respectively) in high throughput situations, as great care is necessary when using 293T-derived cells whose adhesive properties during washing steps are suboptimal.

A key caveat associated with the measurement of neutralizing antibody activity, whether using pseudotype assays or authentic SARS-CoV-2 virions, is that the level of neutralizing activity required to protect against SARS-CoV-2 infection in a natural situation is unknown (Kellam and Barclay, 2020). Moreover, while neutralization assays measure the ability of antibodies to inhibit viral entry, they do not capture features of the antiviral activity of antibodies such as antibody dependent cellular cytotoxicity that may be germane *in vivo* (Bournazos and Ravetch, 2017). Nevertheless, *in vitro* neutralizing activity has long been identified as a correlate of protection against infection in many viral infections (Plotkin, 2010), including coronaviruses (Kellam and Barclay, 2020). As such, we envisage that the techniques described herein might be of significant utility in curtailing the COVID19 pandemic.

## Materials and Methods

### Plasmid constructs

The *env*-inactivated HIV-1 reporter construct pHIV-1_NL4-3_ ΔEnv-NanoLuc was generated from pNL4-3 (Adachi et al., 1986). The human codon-optimized NanoLuc Luciferase reporter gene (*Nluc*, Promega) was inserted in place of nucleotides 1-100 of the *nef*-gene. Thereafter a 940 bp deletion and frameshift was introduced into *env*, immediately 3’ to the *vpu* stop-codon, The pHIV-1_NL_GagPol has previously been described.

The pCCNG/nLuc was construct was derived from pCSGW by inserting a CMV promoter in place of the native SFFV promoter. Thereafter, a NanoLuc-(FMDV2A)-EGFP cassette was inserted 3’ to the CMV promoter.

The rVSVΔG/NG/NanoLuc plasmid was derived from rVSVΔG (Kerafast) (PMID: 20709108). A cassette containing an mNeonGreen/FMDV2A/NanoLuc luciferase cDNA was generated by overlap extension PCR and inserted between the M and L genes, maintaining the intergenic VSV sequences required for gene expression.

Two pSARS-CoV-2 protein expression plasmids containing a C-terminally truncated SARS-CoV-2 S protein (pSARS-CoV-2_Δ19_) were generated. One was derived by insertion of a synthetic human-codon optimized cDNA (Geneart) encoding SARS-CoV-2 S lacking the C-terminal 19 codons into pCR3.1. A second construct was derived from a codon-optimized plasmid (SinoBiological) behaved identically in our assays and were used interchangeably.

To construct a replication competent rVSV/SARS-CoV-2 chimeric virus clone, a codon-optimized cDNA sequence encoding the SARS-CoV-2 spike protein (SinoBiological) but lacking the C-terminal 18 codons was inserted, using Gibson cloning, into a recombinant VSV background that contains GFP immediately upstream of the L (polymerase) following a strategy we previously described for the exchange of VSV-G with HIV-1 Env proteins (Liberatore et al., 2019).

An ACE2 lentivirus expression CS(ACE2)IB vector was constructed by inserting a cDNA encoding an unaltered ACE2 (Addgene:1786) or an catalytically inactive ACE2 mutant (ACE2-H374N&H378N) into the lentivirus expression vector CSIB (Kane et al., 2018). In this vector, expression of the inserted cDNA is driven by a an SSFV promoter and is linked to an IRES-blasticidin.

### Cell lines

HEK-293T cells, HT1080 cells, Huh-7.5 hepatoma cells (*H. sapiens*) and VeroE6 kidney epithelial cells (*Chlorocebus sabaeus*) were cultured in Dulbecco’s Modified Eagle Medium (DMEM) supplemented with 1% nonessential amino acids (NEAA) and 10% fetal bovine serum (FBS) at 37°C and 5% CO_2_. All cell lines have been tested negative for contamination with mycoplasma and were obtained from the ATCC (with the exception of Huh-7.5). Derivatives of 293T and HT1080 cells expressing ACE2 or ACE2* (a catalytically inactive mutant of ACE2) were generated by transducing 293T cells with CSIB(ACE2) or CSIB(ACE2*), respectively. Cells were used as an uncloned bulk populations (designated (B)): 293T/ACE2*(B) and 293T/ACE2(B). Alternatively, single cell clones (cl.) were derived by limiting dilution from the bulk populations and are designated 293T/ACE2*cl.13, 293T/ACE2*cl.21, 293T/ACE2cl.16, 293T/ACE2cl.22 and HT1080/ACE2cl.14

### Two-plasmid-based (HIV/NanoLuc)-SARS-CoV-2 pseudotype particles

To generate (HIV/NanoLuc) SARS-CoV-2 pseudotype particles, 5×10^6^ 293T cells were plated per 10 cm dish in 10ml in growth medium. The following day, 7.5 µg of pHIV-1_NL4-3_ ΔEnv-NanoLuc reporter virus plasmid and 2.5 µg of a SARS-CoV-2 or SARS-CoV plasmid (unless otherwise indicated pSARS-CoV-2-S_Δ19_ was used) were mixed mix thoroughly with 500 µl serum-free DMEM (this represents a molar plasmid ratio of 1:0.55). Then, 44 µl polyethylenimine (PEI, 1 mg/ml) was diluted in 500 µl serum free DMEM and mixed thoroughly.

To generate control virus lacking S, the S expression plasmid was omitted from the transfection and PEI amount was reduced to 30 µl. The diluted DNA and PEI were then mixed thoroughly by pipetting or vortexing, incubated at 20 min at RT and added dropwise to the 293T cells. After 8h or overnight incubation, the transfected cells are washed carefully twice with PBS and incubated in 10ml DMEM++. At 48h after transfection, the 10ml supernatant was harvested, clarified by centrifugation at 300xg for 5 min and passed through a 0.22 µm pore size PVDF syringe filter (e.g. SLGVR33RS, Millipore), aliquoted and frozen at -80°C

### Three-plasmid-based (HIV-1/NG/NanoLuc)-SARS-CoV-2 pseudotype particles

To generate (HIV-1/NanoLuc2AEGFP)-SARS-CoV-2 particles, 3 plasmids were used, with the reporter vector (pCCNanoLuc2AEGFP) and HIV-1 structural/regulatory proteins (pHIV_NL_GagPol) provided by separate plasmids. Specifically, 293T cells were transfected as described above, with 7 µg pHIV_NL_GagPol, 7 µg pCCNanoLuc2AEGFP and 2.5 µg 2.5 µg of a SARS-CoV-2 or SARS-CoV plasmid (unless otherwise indicated pSARS-CoV-2-S_Δ19_ was used, at a molar plasmid ratio of 1:1:0.45) using 66 µl PEI. To generate control virus lacking the S expression plasmid was omitted from the transfection and PEI amount was reduced to 56µl. At 48h after transfection, the 10ml supernatant was harvested, clarified, filtered and stored as described above

### Recombinant VSVΔG-based (VSV/NG/NanoLuc)-SARS-CoV-2 pseudotype particles

To generate (VSV/NG/NanoLuc)-SARS-CoV-2 pseudotype particles 293T cells were plated at 1×10^6^ cells/well in 6-well plates. The following day cells were rinsed with with serum free medium (SFM) and infected with recombinant T7-expressing vaccinia virus (vTF7-3) in SFM at MOI of ∼5 for 30-45 min, gently rocking the plate every 10-15 minutes. Thereafter, cells were washed with DMEM and 1.5 ml DMEM++ added per well. Next a mixture of plasmids encoding the rVSV antigenome; rVSVΔG/NG/NanoLuc, (500ng) and rescue plasmids pBS-N (300ng) pBS-P (500ng) pBS-L (100ng) pBS-G (800ng) were mixed with 5.5 ul PLUS reagent in 100ul Opti-MEM. Then, 9ul Lipofectamine LTX was added to 125ul Opti-MEM, and the diluted plasmid DNA and Lipofectamine LTX mixed and incubated for 20 minutes prior to addition to the vTF7-3-infected cells. The growth medium was replaced the following morning. At ∼24h post transfection the supernatant was collected, filtered through a 0.1μm filter and used to infect VSV-G expressing cells for amplification.

To amplify rescued rVSVΔG/NG/NanoLuc. 5×10^6^ 293T cells were plated per 10 cm dish in 10ml in growth medium or 1.2×10^7^ 293T cells were plated in 15cm dishes. The following day, cells were transfected with 5μg (10cm dish) or 12.5μg (15cm dish) of pCMV-VSV-G expression plasmid, using PEI. The following day, the transfected cells were infected with the rescued virus and 16h later the supernatant was collected centrifuged at 350g to clarify and filtered through a 0.22μm filters.

To prepare stocks of (VSV/NG/NanoLuc)-SARS-CoV-2 pseudotype particles, 293T cells were plated at 1.2×10^7^ 293T cells were plated in 15cm dishes, and transfected the following day with 12.5μg pSARS-CoV2_Δ19_. The next day the transfected cells were infected with the above described VSV-G complemented rVSVΔG/NG/NanoLuc virus at an MOI of 1. At 16h later the supernatant was collected, centrifuged at 350xg to clarify and filtered through a 0.22μm filter. Next the filtered supernatant was layered on top of a 20% sucrose cushion and centrifuged at 25,000 rpm for 1.5h in a SW32 Ti rotor in a Beckman Optima XE-90 Ultracentrifuge. Alternatively, virions were concentrated using Lenti-X-Concentrator (Takarabio). The pelleted virus was resuspended in DMEM aliquoted and stored at -80°C. Prior to infection of target cells, the viral stock was incubated with 20% I1 hybridoma (anti-VSV-G) supernatant (ATCC CRL-2700) for 1h at 37°C to neutralize contaminating rVSVΔG/NG/NanoLuc/VSV-G particles.

### Replication competent VSV/SARS-CoV-2 chimera

To recover the infectious rVSV/SARS-CoV-2/GFP chimeric virus, 293T cells were plated in 6-well plates infected with vTF7-3 and transfected with a mixture of plasmids encoding the rVSV/SARS-CoV-2/GFP (500ng) and rescue plasmids including pBS-G, as described above for VSVΔG pseudotype particles. At ∼24h post transfection the supernatant was collected, filtered through a 0.1μm filter to remove vaccina virus and used to infect 293T cells transfected with the pCMV-VSV-G expression plasmid for amplification of the rVSV/SARS-CoV-2/GFP population. Thereafter, the complemented virus was used to infect 293T/ACE2(B) cells in 25cm^2^ flasks in the absence of the complementing VSV-G protein, and the virus population passaged, as described in the Results section. To isolate adapted variants, the viral supernatant was serially diluted and aliquots of each dilution used to inoculate 12 wells of a 96-well plate containing 293T/ACE2(B) cells. Virus was harvested from wells that contained individual green fluorescent plaques (signifying infection by single viruses). Two plaque purified viruses that were investigated further were designated rVSV/SARS-CoV-2/GFP_1D7_ and rVSV/SARS-CoV-2/GFP_2E1_. RNA was isolated from cultures infected with rVSV/SARS-CoV-2/GFP_1D7_ and rVSV/SARS-CoV-2/GFP_2E1_ using the QIAamp Viral RNA mini kit (Qiagen) and cDNA synthesis was performed using SuperScript III using hexamers (ThermoFisher). Sequences encoding S and flanking regions were PCR amplified and sequenced (Genewiz).

### Infectivity assays

To measure the infectivity of pseudotyped or chimeric viral particles, viral stocks were serially diluted and 100 µl of each dilution added to target cells plated at 1×10^4^ cells/well in 100 µl medium in 96-well plates the previous day. Cells were then cultured for 48h hours (HIV-1 pseudoviruses) or 16h (VSV pseudoviruses or replication competent rVSV/SARs-CoV-2) unless otherwise indicated, and then photographed or harvested for flow cytometry or NanoLuc luciferase assays.

### Neutralization assays

To measure neutralizing antibody activity in plasma, serial dilutions of plasma from COVID19 patients and healthy donors a 1:12.5 initial dilution were five-fold serial diluted in 96-well plates over 7-8 dilutions. To measure neutralization activity of monoclonal antibodies, a 40 µg/ml initial dilution was four-fold serially diluted over 11 dilutions. Thereafter, a 55 µl aliquot of serially diluted plasma, monoclonal antibody or decoy was incubated with a 55 µl aliquot of HIV-1 (2-plasmid), HIV-1 (3-plasmid) or VSV-based SARS-CoV-2 pseudovirus or rVSV/SARS-CoV-2/GFP containing approximately 1×10^3^ infectious units for 1h at 37°C in a 96-well plate. Thereafter, 100 µl of the mixture was added to target cells plated at 1×10^4^ cells/well in 100 µl medium in 96-well plates the previous day. Thus, the final starting dilutions were 1:50 for plasma and 10 µg/ml for monoclonal antibodies. Cells were then cultured for 48h hours (HIV-1 pseudoviruses) or 16h (VSV pseudovirus and rVSV/SARS-CoV-2) unless otherwise indicated. Thereafter cells were photographed or harvested for flow cytometry or NanoLuc luciferase assays

### Reporter gene assays

For the NanoLuc luciferase assays, cells were washed twice, carefully, with PBS and lysed with 50 µl/well of Luciferase Cell Culture Lysis reagent (Promega). NanoLuc Luciferase activity in lysates was measured using the Nano-Glo Luciferase Assay System (Promega). Specifically, 25 µl of substrate in NanoGlo buffer was mixed with 25 µl cell lysate in black flat bottom plates and incubated for 5 min at RT. NanoLuc luciferase activity was measured using a Modulus II Microplate Multimode reader (Turner BioSystem) or a Glowmax Navigator luminometer (Promega), using 0.1s integration time. Relative luminescence units (RLU) obtained were normalized to those derived from cells infected with SARS-CoV-2 pseudovirus in the absence of plasma/antibodies.

To record GFP+ cells, 96-well plates were photographed using and EVOS M7000 automated microscope. Alternatively, cells were trypsinized, fixed with 2% paraformaldehyde, washed and enumerated using an Attune NxT flow cytometer equipped with a 96-well autosampler.

The half maximal inhibitory concentration for plasma (NT_50_), antibodies (IC_50_) was determined using 4-parameter nonlinear regression curve fit to raw infectivity data measured as relative light units, or as the percentage of infected cells (GraphPad Prism). The top values were unconstrained, the bottom values were set to zero.

### SARS-CoV-2 virus stocks and titration

SARS-CoV-2, strain USA-WA1/2020, was obtained from BEI Resources and amplified in VeroE6 cells at 33°C. Viral titers were measured on VeroE6 cells by standard plaque assay (PA). Briefly, 500µl of serial 10-fold virus dilutions in Opti-MEM were used to infect 4×10^5^ cells/well seeded the previous day in 6-well plates. After 90 min adsorption, the virus inoculum was removed, and cells were overlayed with DMEM containing 10% FBS with 1.2% microcrystalline cellulose (Avicel). Cells were incubated for five days at 33°C, followed by fixation with 3.5% formaldehyde and crystal violet staining for plaque enumeration. All experiments were performed in a BSL-3 laboratory.

### SARS-CoV-2 Neutralization Assay

The day prior to infection VeroE6 cells were seeded at 1×10^4^ cells/well into 96-well plates. Plasma samples and antibodies were serial diluted in BA-1, consisting of medium 199 (Lonza, Inc.) supplemented with 1% bovine serum albumin (BSA) and 1x penicillin/streptomycin. Next, the diluted samples were mixed with a constant amount of SARS-CoV-2 and incubated for 60 min at 37 °C. The plasma/antibody-virus-mix was then directly applied to VeroE6 cells (MOI of ∼0.1 PFU/cell; n=3) and incubated for 18-20 h at 37 °C Cells were subsequently fixed by adding an equal volume of 7% formaldehyde to the wells, followed by permeabilization with 0.1% Triton X-100 for 10 min. After extensive washing, cells were incubated for 1 h at RT with blocking solution of 5% goat serum in PBS (catalog no. 005–000-121; Jackson ImmunoResearch). A rabbit polyclonal anti-SARS-CoV-2 nucleocapsid antibody (catalog no. GTX135357; GeneTex) was added to the cells at 1:500 dilution in blocking solution and incubated at 4 °C overnight. Alternatively, J2, a mouse monoclonal anti-dsRNA antibody (catalog no. 10010500; Scicons) was added to the cells under similar conditions to detect virus infected cells. Goat anti-rabbit AlexaFluor 594 (catalog no. A-11012; Life Technologies) and goat anti-mouse AlexaFluor 488 (catalog no. A-11001; Life Technologies) were used as a secondary antibodies at a dilution of 1:2,000. Nuclei were stained with Hoechst 33342 (catalog no. 62249; Thermo Scientific) at a 1:1,000 dilution. Images were acquired with a fluorescence microscope and analyzed using ImageXpress Micro XLS (Molecular Devices, Sunnyvale, CA).

### Human plasma samples and monoclonal antibodies

The human plasma and monoclonal antibodies used in this study were previously reported (Robbiani et al., 2020). The human samples were obtained at the Rockefeller University Hospital under protocols approved by the University’s Institutional Review Board.

## Supplemental material

### Supplemental figures

**Fig. S1. Generation of and HIV-1 pseudotype infection of ACE2-expressing cell lines**.

**Fig. S2. Variables determining HIV-1 pseudotype infection signal**.

**Fig. S3. rVSVΔG/NG-NanoLuc pseudotyped virus infection**.

**Fig. S4. Examples of neutralization of HIV-1 and VSV pseudotyped virus particles by monoclonal antibodies targeting SARS-CoV-2 S**.

## Author Contributions

PDB TH MCN CMR and DR conceived and supervised the studies. FS, YW, FM and EB built recombinant viral plasmids. YW and MR developed ACE2-expressing cell lines. FS and YW developed and performed the VSV pseudotype and chimeric virus assays. FM and JCCL developed and performed the HIV-1 pseudotype assays with assistance from PM. HHH and EM developed and performed the SARS-CoV-2 neutralization assays. MC and CG provided clinical samples, MA, AC and ZW discovered and cloned monoclonal antibodies that were produced and purified by AG and MC. PDB and TH wrote the manuscript with input from other authors.

## Acknowledgements

This work was supported by NIH grants P01AI138398-S1, 2U19AI111825 (to M.C.N. and C.M.R), R01AI091707-10S1 (to C.M.R.) R01AI078788 (to T.H.) R37AI64003 (to P.D.B.) and by the George Mason University Fast Grant (to D.F.R. and to C.M.R.) and the European ATAC consortium EC101003650 (to D.F.R) and the G. Harold and Leila Y. Mathers Charitable Foundation (to C.M.R.). C.G. was supported by the Robert S. Wennett Post-Doctoral Fellowship, in part by the National Center for Advancing Translational Sciences (National Institutes of Health Clinical and Translational Science Award program, grant UL1 TR001866), and by the Shapiro-Silverberg Fund for the Advancement of Translational Research. P.D.B. and M.C.N. are Howard Hughes Medical Institute Investigators.

---

**Fig. S1.**
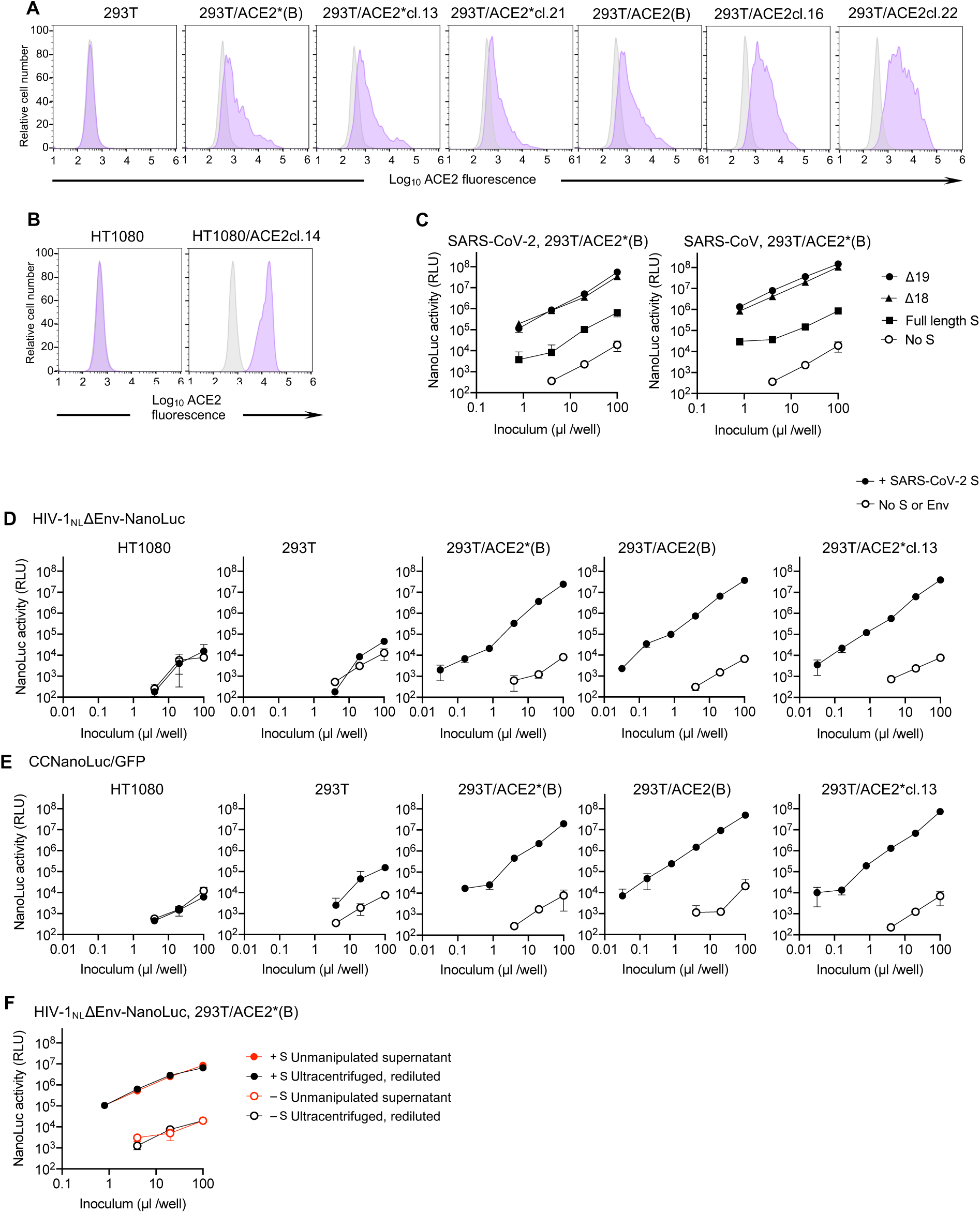
Generation of and HIV-1 pseudotype infection of ACE2-expressing cell lines. **A**. 293T cells were stably transduced with a lentivirus vector CSIB, expressing either wild type ACE2 or catalytically active mutant ACE2*. Following selection, cells were used as uncloned bulk populations (B) or single cell clones were isolated. Flow cytometry histograms show staining with an antibody against huACE2 (purple) or an isotype control (grey). **B**. HT1080 cells were stably transduced as in A and a single cell clone used throughout this study is shown, stained as in A. **C**. Infectivity of CCNanoLuc/GFP viruses, pseudotyped with either full length or C-terminally truncated SARS-CoV and SARS-CoV-2 S proteins on 293T/ACE2*(B) cells. Virus particles generated in the absence of an S protein (No S) were used as background controls. Infectivity was quantified by measuring NanoLuc luciferase activity (RLU). Average and standard deviation from two technical replicates is shown. **D**. Infectivity of HIV-1_NL_ΔEnv-NanoLuc in the various cell lines. Virus generated in the absence of S is used as a background control and infectivity was quantified by measuring NanoLuc luciferase activity (RLU). Average and standard deviation from two technical replicates is shown. **E**. Same as D except that CCNanoLuc/GFP virus was used **F**. Effect of virus ultracentrifugation on the infectivity of HIV-1-based pseudotyped virus particles. 293T/ACE2*(B) cells were infected with equivalent doses of unconcentrated HIV-1_NL_ΔEnv-NanoLuc, or the same virus that had be pelleted through 20% sucrose and then diluted to the original volume.

**Fig. S2.**
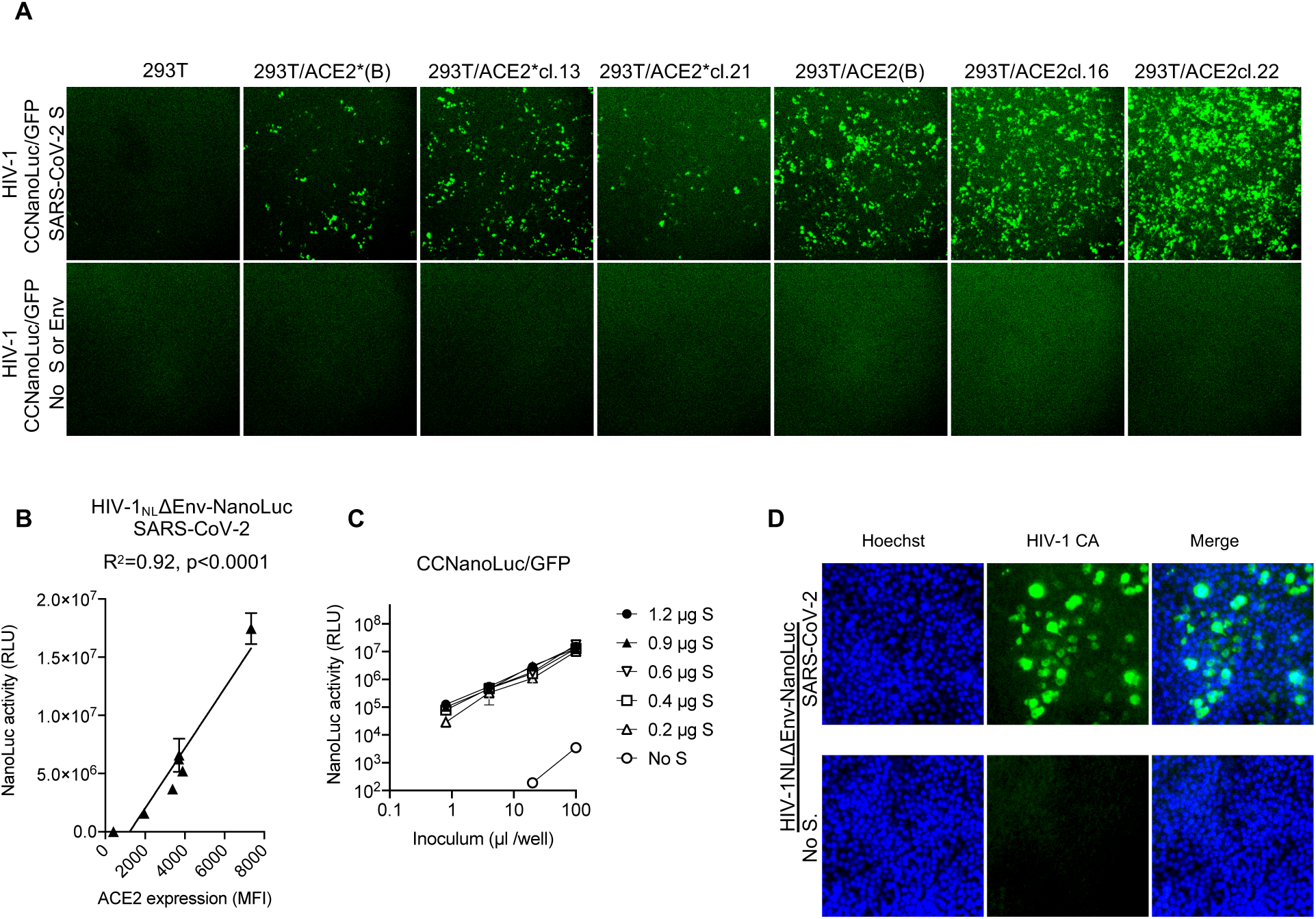
Variables determining HIV-1 pseudotype infection signal. **A**. Images (GFP) of confluent monolayers of the indicated cell lines after infection with equivalent amounts of CCNanoLuc/GFP pseudotyped with SARS-CoV-2 SΔ19 or no S protein, as indicated. **B**. Relationship between NanoLuc luciferase activity (RLU) and ACE2 cell surface expression levels (quantified by flow cytometry, Fig S1A) following infection the cell lines depicted in Fig. S1A with HIV-1_NL_ΔEnv-NanoLuc pseudotyped virus. **C**. Infectivity of CCNanoLuc/GFP pseudotyped virus generated by cotransfection with the indicated amounts of SARS-CoV-2 SΔ19 expression plasmid. **D**. Quantification of infectivity of HIV-1_NL4-3_ ΔEnv-NanoLuc pseudotype infection by immunostaining of 293T/ACE2(B) target cells with antibodies against the HIV-1 capsid (CA) protein. Nuclei were visualized by Hoechst staining.

**Fig. S3.**
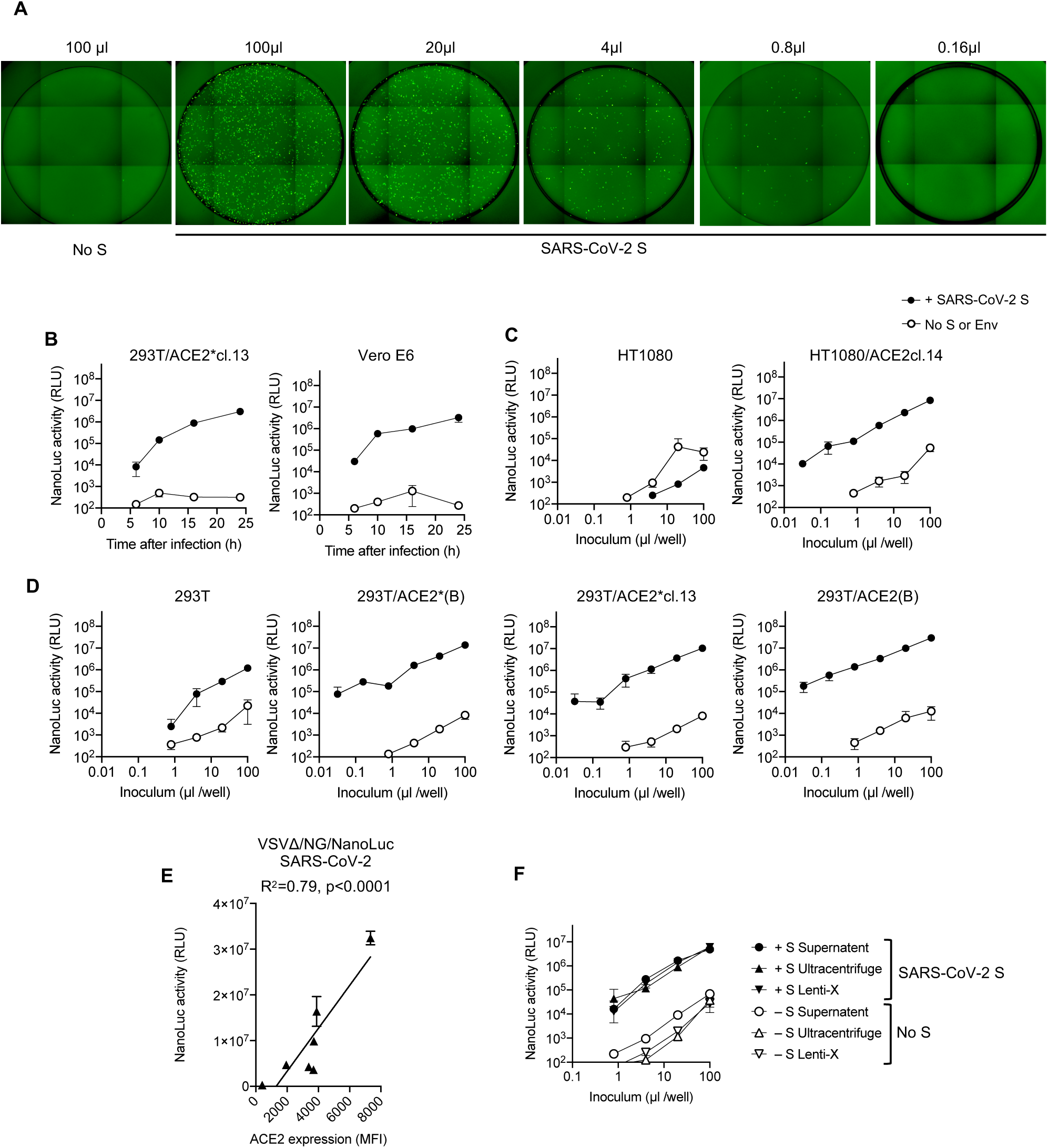
rVSVΔG/NG-NanoLuc pseudotyped virus infection. **A**. Infection of Huh7.5 cells with the indicated volumes of rVSVΔG/NG-NanoLuc pseudotyped virus. Images of the entire well of 96-well plates are shown. **B**. Infectivity of rVSVΔG/NG-NanoLuc pseudotyped with SARS-CoV-2 SΔ19 or no S (background control) on the indicated cell lines. Infectivity was quantified at the indicated times post-inoculation by measuring NanoLuc luciferase levels (RLU). **C and D** HT1080-derived cell lines (C) or 293T-derived cell lines (D) were infected with varying amounts of rVSVΔG/NG-NanoLuc pseudotyped with SARS-CoV-2 SΔ19 or no S (background control) and NanoLuc luciferase levels were measured at 16h after infection. **E**. Relationship between NanoLuc luciferase activity (RLU) and ACE2 cell surface expression levels (quantified by flow cytometry, Fig S1A) following infection the cell lines depicted in Fig. S1A with rVSVΔG/NG-NanoLuc. **F**. Effect of virus concentration on the infectivity of rVSVΔG/NG-NanoLuc pseudotyped virus. Huh7.5 cells were infected with equivalent doses of either unmanipulated virus-containing supernatant or virions that had been pelleted by ultracentifugation or using Lenti-X and diluted to the original volume.

**Fig. S4.**
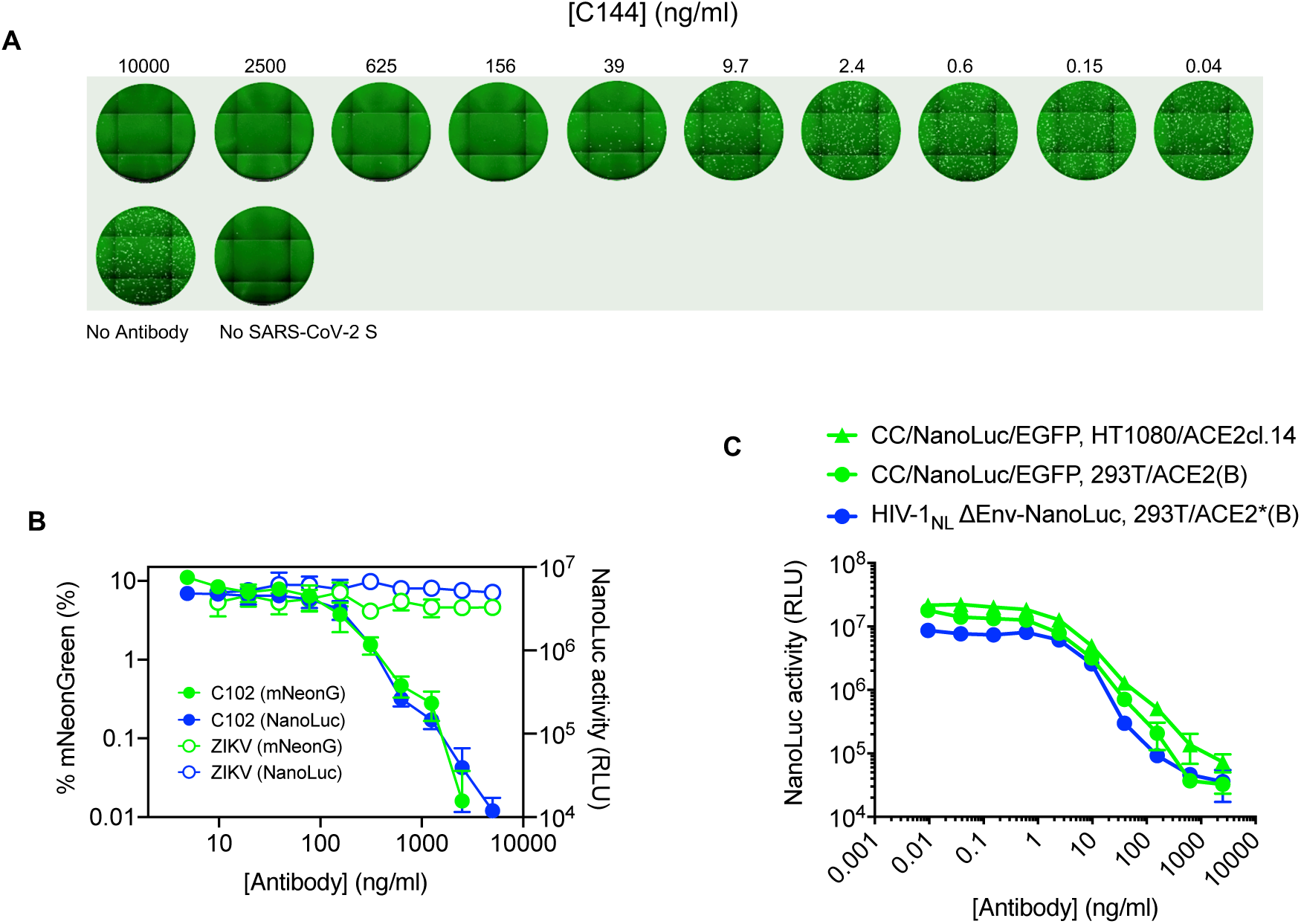
Examples of neutralization of HIV-1 and VSV pseudotyped virus particles by monoclonal antibodies targeting SARS-CoV-2 S. **A**. Images of Huh7.5 cells following infection with rVSVΔG/NG-NanoLuc pseudotyped virus (∼10^3^ IU/well) in the presence of the indicated concentrations of a human monoclonal antibody (C144) targeting SARS-CoV-2 S RBD. **B**. Quantification of rVSVΔG/NG-NanoLuc pseudotyped virus infection (measured by flow cytometry (% mNeonGreen positive cells, green) or by NanoLuc luciferase activity (RLU, blue) in the presence of the indicated concentrations of a human monoclonal antibody (C102) targeting SARS-CoV-2 S RBD, or a control monoclonal antibody against the Zika virus envelope glycoprotein. **C**. Quantification of HIV-1_NL_ΔEnv-NanoLuc or CCNanoLuc/GFP pseudotyped virus infection on the indicated cell lines in the presence of the indicated concentrations of a human monoclonal antibody (C121) targeting SARS-CoV-2 S RBD Infectivity was quantified by measuring NanoLuc luciferase levels (RLU).

